# Social media reveals consistently disproportionate tourism pressure on a threatened marine vertebrate

**DOI:** 10.1101/2020.06.17.157214

**Authors:** Kostas Papafitsoros, Aliki Panagopoulou, Gail Schofield

## Abstract

Establishing how wildlife viewing pressure is distributed across individual animals within a population can inform the management of this activity, and ensure targeted individuals or groups are sufficiently protected. Here, we used social media data to quantify whether tourism pressure varies in a loggerhead sea turtle (Caretta caretta) population and elucidate potential implications. Laganas Bay (Zakynthos, Greece) supports both breeding (migratory, and hence transient) and foraging (resident) turtles, with turtle viewing representing a major component of the tourism industry. Social media entries spanning two seasons (April to November, 2018 and 2019) were evaluated, and turtles were identified via photo-identification. For both years, 1684 and 2105 entries of 139 and 122 unique turtles were obtained from viewings, respectively (boats and underwater combined). However, while residents represented less than one-third of uniquely identified turtles, they represented 81.9% and 87.9% of all entries. Even when the seasonal breeding population was present (May to July), residents represented more than 60% entries. Of note, the same small number of resident turtles (<10), mostly males, were consistently viewed in both years; however, different individuals were targeted by boats versus underwater. Thus, turtles appear to use and remain in the area despite high viewing intensity, possibly indicating low disturbance. However, photo-identification records revealed a high risk of propeller and boat strike to residents (30%) leading to trauma and mortality. To reduce this threat and ease viewing pressure, we recommend the compulsory use of propeller guards for all boats and the creation of temporary “refuge” zones for resident animals at viewing hotspots, with these suggestions likely being relevant for other wildlife with similar population dynamics. In conclusion, social media represents a useful tool for monitoring individuals at a population scale, evaluating the pressure under which they are placed, and providing sufficient data to refine wildlife viewing guidelines and/or zoning.

## Introduction

Ecotourism focusing on wildlife is a growing industry promoting sustainable conservation and community development through the economic benefits it generates (Kals et al., 1999; Balmford et al., 2015). Over the last 20 years, this industry has expanded from targeting primarily terrestrial species to include a wealth of marine fauna, including marine mammals (e.g. Holcomb et al., 2009; Christiansen et al., 2014), sea turtles (e.g. Schofield et al., 2015), sharks (e.g. Cisneros-Montemayor et al., 2013) and rays (e.g. Semeniuk et al., 2009; Barr and Abelson, 2019). Parallel research on how this industry impacts wildlife has led to contradictory results, even within the same species, and is thought to be dependent on the environment and activity of the animal (Gill et al., 2001; Williams et al., 2009; New et al., 2013). Yet, it is important to quantify how pressure derived from viewing varies both spatially and temporally within a population to identify individuals or groups (e.g. immature versus adult animals, residents versus non-residents, or a particular sex) that are disproportionately targeted, and the implications on survival and fitness (Holcomb et al., 2009; Semeniuk et al., 2009; Christiansen and Lusseau, 2014). For instance, resident animals tend to be subject to higher viewing pressure than seasonal migrants, due to operators targeting sites known to be regularly used by these individuals to guarantee viewings (Christiansen and Lusseau, 2014; Schofield et al., 2015; Barr and Abelson, 2019).

Wildlife watching is intrinsically linked with taking photographs, which creates opportunities to collect important baseline data for ecological research at unprecedented spatial and temporal scales (Dickinson et al., 2012; Toivonen et al., 2019). Photo-identification is a minimally invasive approach used to identify unique individuals in a given population, and has become widely adopted by the scientific community, from research on seadragons to giraffes (Martin-Smith, 2011; Halloran et al., 2015). This approach is also providing opportunities for the public to become directly involved in projects as citizen scientists, by contributing their photographs and increasing the quantity of available data (Holmberg et al., 2008). The ubiquity of mobile phones and image sharing (photographs and videos) on social media presents another emerging image-based resource for use in conservation science (Di Minin et al., 2015; Toivonen et al., 2019). This medium has great potential to provide qualitative and quantitative information about various human-wildlife interactions, including recreational fishing (e.g. Martin et al., 2014; Giovos et al., 2018), wildlife trade (e.g. Di Minin et al., 2019), and wildlife distributions (e.g. Baumbach et al., 2019; Casale et al., 2020).

Sea turtles are globally threatened marine vertebrates that are subjected to wildlife viewing both on land (during nesting on beaches, e.g. Tisdell and Wilson, 2005) and in coastal waters (Schofield et al., 2015). Several studies have already attempted to infer the behavioral responses of turtles to the presence of swimmers/snorkelers (e.g. Griffin et al., 2017; Hayes et al., 2017). However, these studies are typically conducted at small scales, with logistical issues limiting the quantification of encounters at a large scale in space and time. It has been previously demonstrated that the intensity of viewing sea turtles in the marine environment changes with the number of animals available for viewing, affecting operator strategies approach and level of compliance with existing guidelines, with implications for species that aggregate in seasonal populations for breeding or foraging (Schofield et al., 2015). This staggered arrival or departure of individuals likely leads to certain individuals or groups of individuals being disproportionately targeted; however, the tools to evaluate this phenomenon were not available until recently. Here, we aimed to address this knowledge gap at an important breeding site for loggerhead sea turtles in the Mediterranean by combining data derived from social media and photo-identification techniques to evaluate whether wildlife viewing activities disproportionately target certain individuals at a site used by both resident and migrant individuals. Laganas Bay on Zakynthos, Greece, supports a seasonal breeding aggregation of over 200 loggerhead sea turtles (*Caretta caretta*) (from April to early August of each year), but also a small population of resident male and immature turtles (Schofield et al., 2015; Dujon et al., 2018). This site was ideal for our study because it is also a popular summer tourist destination in the Mediterranean attracting over 850,000 visitors between late-April and late-October, and it supports wildlife-watching activities. We hypothesized that resident turtles would be subject to greater viewing pressure during the late part of the breeding season, when the numbers of breeding turtles frequenting the site decline. The results of this study are expected to demonstrate the utility of social media in quantifying viewing pressure on certain sub-groups in a given population and providing evidence to revise viewing guidelines with operator compliance.

## Methods

### Study site and species

Laganas Bay, on the Greek island of Zakynthos (Figure 1a; 37*°*43^*′*^ N, 20*°*52^*′*^ E) hosts a major loggerhead sea turtle nesting site, which is protected within the framework of the National Marine Park of Zakynthos (NMPZ) established 1999 (Figure 1a; Margaritoulis, 2005). The park encompasses a terrestrial zone including the nesting sites, and a marine area divided into three zones to ensure protection of turtles during the entire nesting season. The maritime zoning of the NMPZ was precautionary based, reflecting the intensity of nesting activity on the six nesting beaches, rather than the distribution of turtles in the water. Consequently, no boating activity is allowed in Zone A, where most nesting activity on beaches occurs; however, boating at 6 knots is permitted in Zones B and C where over 70% of turtles aggregate in the marine area, resulting in a large overlap in the distribution of tourists and turtles in the nearshore coastal area (Zbinden et al., 2007; Schofield et al., 2013). Wildlife-watching activities were initiated as an alternative source of income to water-sports for the local community in Laganas Bay in the mid-1990s, with the NMPZ providing guidelines aiming to minimize pressure on turtles observed. Since 2018, these guidelines include a legislative act issued by the local coast guard specifying minimum observation distances, durations and number of vessels observing an animal. This activity is currently estimated to service 180000 tourists over 9,000 trips per year, generating an annual revenue of over 2.7 million euros (data adjusted from Schofield et al., 2015, setting 20 tourists per trip, 15 euros per ticket); however, because 100s of turtles aggregate within 500 m of shore (Schofield et al., 2017), it is possible to view them by wading, snorkeling or swimming and from privately used vessels (pedalloes, kayaks, private hire boats) (Schofield et al., 2015). Snorkelling with sea turtles is not regulated by law and it is allowed in all zones of the NMPZ.

**Figure 1.**
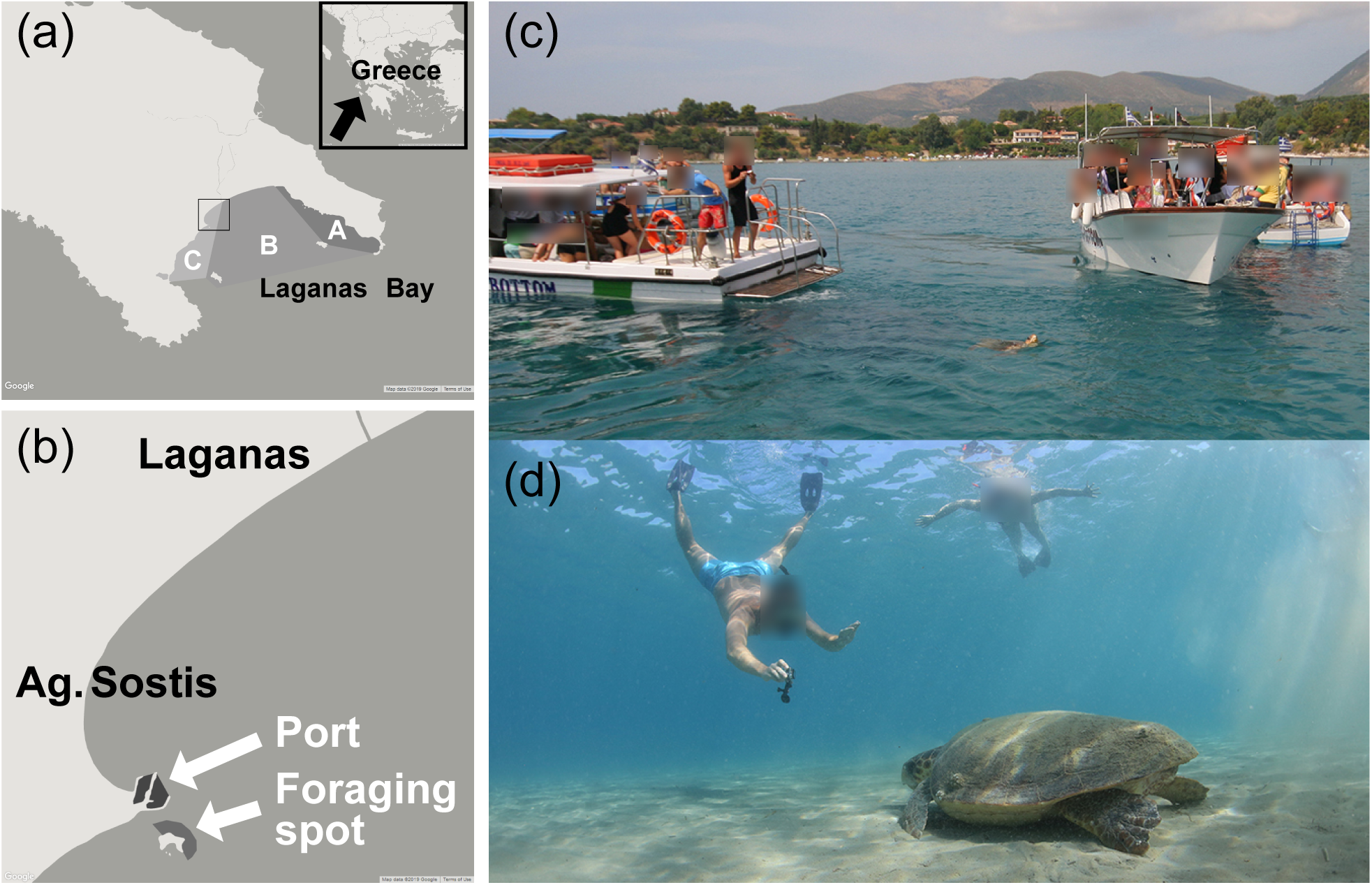
(a) Southern part of Zakynthos Island (Greece) showing Laganas Bay and the maritime zoning of the National Marine Park of Zakynthos. Black box represents the area where 85% of 1034 social media entries, for which the area could be detected, were obtained (representing 27% out of the total 3789 entries) including (b) Agios Sostis Port (labeled Port) and a foraging spot used by resident turtles. Photographs of (c) wildlife watching vessels observing a turtle and (d) a snorkeler photographing a turtle underwater. Photograph credits: Kostas Papafitsoros.

The loggerhead sea turtle population of the Mediterranean is currently listed as Least Concern in the IUCN Red List of Threatened Species, with its current status regarded to be the result of long-term conservation efforts and it is therefore considered conservation dependent (Casale, 2015). Adult turtles migrate into Laganas Bay to mate as early as March, with most adult males departing in late May, while females typically remain to nest from mid-May to early August, at which point they depart for foraging grounds up to 1000 km distant (Schofield et al., 2013; Schofield et al., 2020). Direct counts of turtles using UAVs indicated the presence of up to 90 unique males and 240 unique females in the breeding area in one season (Schofield et al., 2017), which fell within the range predicted for previous years based on nest counts by (Schofield et al., 2015). Tracking of male and female sea turtles from Zakynthos (n = 20 females, Zbinden et al., 2011; n = 55 males and females, Schofield et al., 2020) has demonstrated that females migrate to other areas of the Mediterranean following breeding, whereas around 30% of males remain resident. This information has been corroborated with long-term photo-identification records of over 1000 unique adult male and females, as well as immature turtles (Schofield et al., 2008; Schofield et al., 2020). Residency is defined as turtles continuing to frequent the area from June onwards (for males) and after females have departed in early August (for females), and at any time for immature turtles (Schofield et al., 2013). Around 40 resident male and immature turtles have been documented through the photo-identification database (Kostas Papafitsoros-*personal observations*).

### Collection of social media records

We used Instagram as the social media platform for this study. While other platforms exist (e.g. Facebook, Flickr, Youtube), we selected Instagram because it has a convenient search method, via the use of hashtags (“#”). We searched for social media videos and photographs (termed entries) from 1 April to 30 November for 2018 and 2019, with this period encompassing the entire tourist season (end of April to end of October).

We searched Instagram using popular searchterms related to our study site, as well as common associated locations and common misspellings of these terms (see Supplementary material for a full list). Hashtags (“#”) and searchable locations in Instagram were used as convenient search frameworks. Searches were completed at least once a week to obtain information in real-time, with retrospective searches being used to confirm no entries were missed. We rejected entries that clearly belonged to previous seasons (e.g. known turtles that died in previous years) and entries used as advertisements by tour agencies. For each entry included in our analyses, we recorded the date uploaded, and date recorded, if available from the caption. For 74 and 97 entries in 2018 and 2019, respectively, the date the images were acquired was provided, of which 63 and 77 (85.1% and 79.4%) were uploaded on the same date or the day after. Overall, the discrepancy between the time an image was obtained to uploading was a mean days (SD=9.4) and 3.4 days (SD=9.1) for 2018 and 2019. We also recorded whether the entries were taken underwater (i.e. usually by people snorkeling/swimming), above water (i.e. from a boat when the turtle was breathing) or of turtles in the water from land (usually the Agios Sostis Port area, Figure 1). For images from boats, we did not differentiate whether they were taken during an organized wildlife tour or a privately hired boat or pedalo. We did not use any personal information of people who uploaded images. For each entry, we manually compared the turtle image against those in our existing 20-year photo-identification database, containing over 1000 unique individuals (Schofield et al., 2020). Photo-identification followed a validated approach using facial scute patterns (Figure 2; Schofield et al., 2008). Only images where facial scutes could be distinguished were used for our analyses (see Supplementary figure 1). The entries were finally classified depending on the location (i.e. underwater, from a boat, or from land). Turtles from the social media entries were only classified as new individuals in the database when the quality of the entry was high.

**Figure 2.**
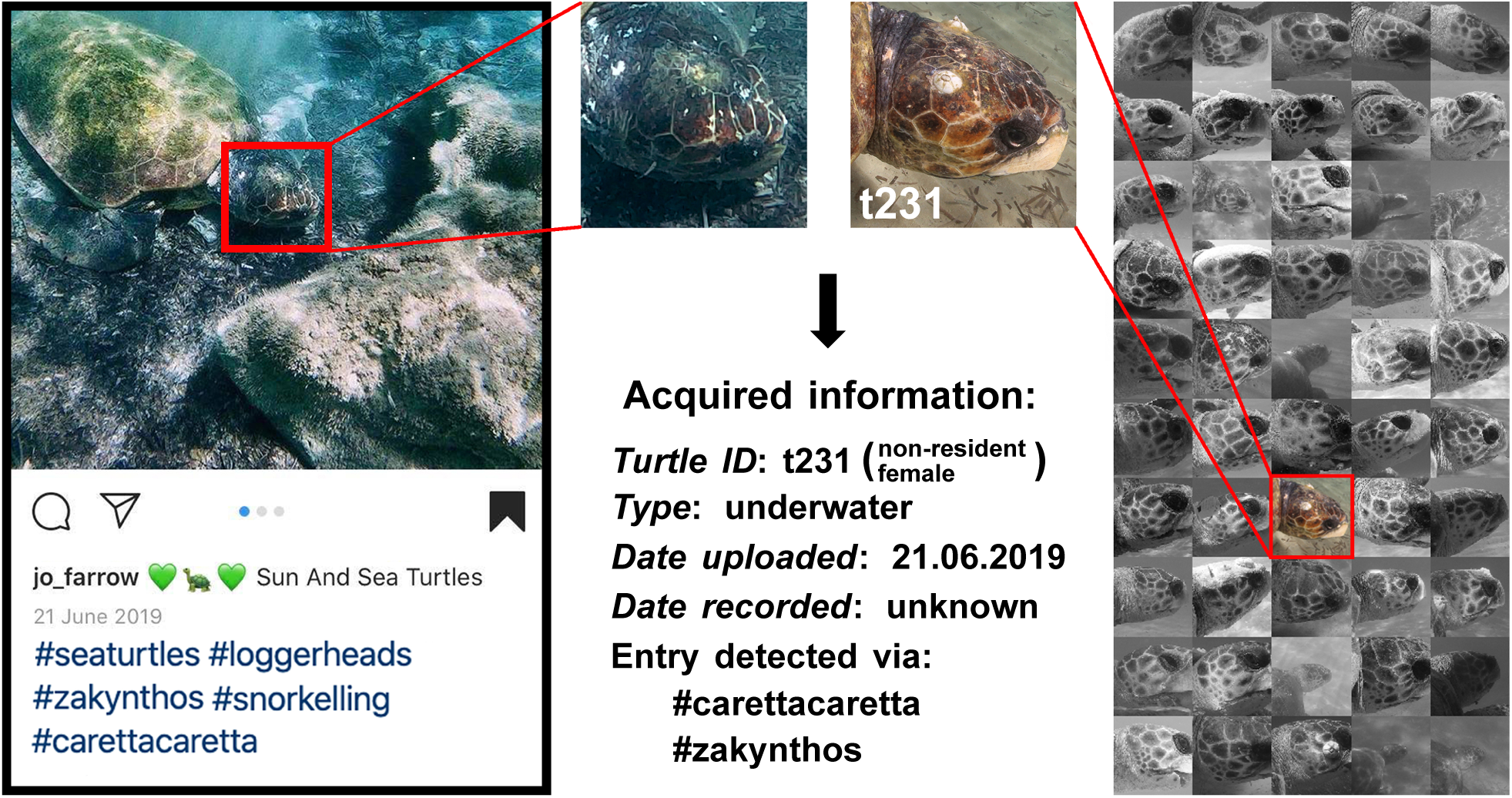
Workflow for data extraction from social media entries. Photograph credits: Jo Farrow (left - used with permission), Kostas Papafitsoros (right).

### Risk of trauma

To assess risk of boat induced trauma, we searched all records of resident turtles present in 2018 and 2019 in our photo-identification database for evidence of damage to the carapace. We first recorded the last date that individuals had no trauma or no new trauma and the date on which a new trauma was detected, to confirm that trauma occurred during the proximate season.

Data on the number of visitors arriving on Zakynthos 2018 and 2019 were obtained from the Hellenic Civil Aviation Authority (CAA) (http://www.ypa.gr). We used MATLAB to visualize the data and to perform descriptive statistics (means, standard deviations, calculation of Pearson correlation coefficient).

## Results

### Instagram records and tourist numbers

We identified 1684 and 2105 (total: 3789) Instagram entries between April and November 2018 and 2019, respectively. In both years, the number of entries peaked in July and August; these two months combined accounted for >52% of entries in each year; Figure 3; Supplementary Table 1). In both years, most entries were taken from a boat (>65%), followed by underwater (>20%) and land (about 6%) (Supplementary Table 2). Because very few individuals were documented from land, we did not evaluate this component further. The monthly percentage of entries was strongly correlated with the monthly number of airport arrivals (Pearson correlation coefficient p = 0.9710 for 2018 and p = 0.9362 for 2019; Figure 3b).

**Figure 3.**
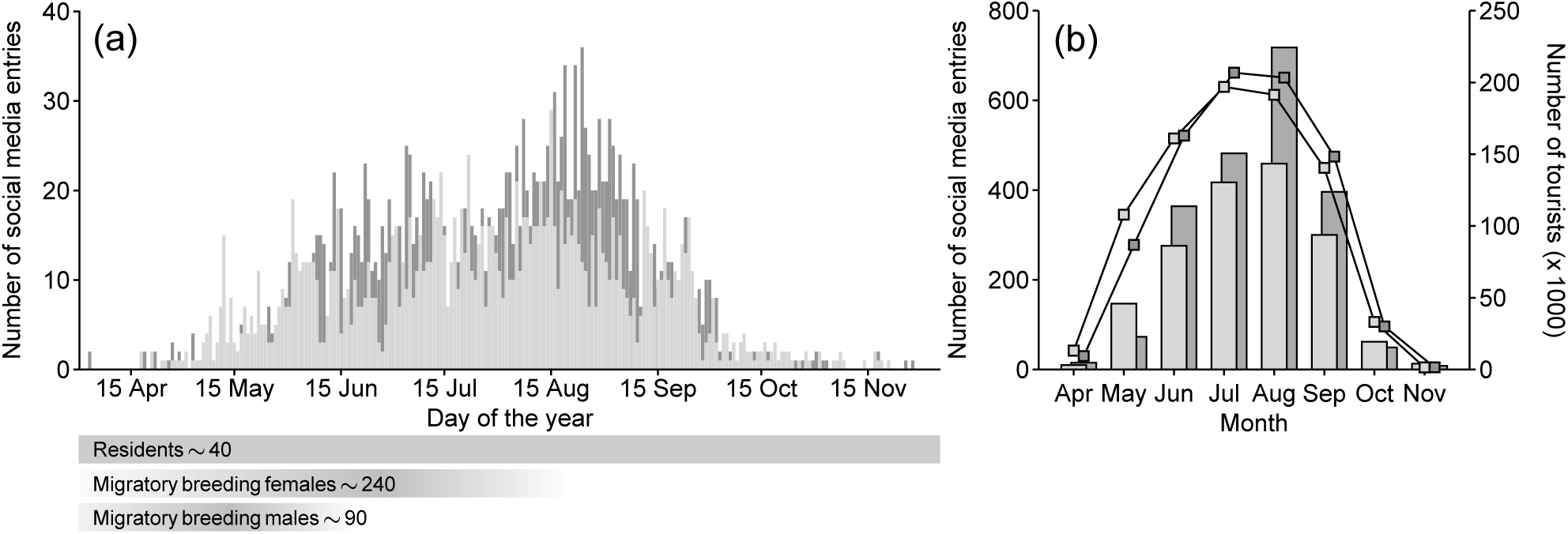
Number of social media entries (bars) (a) daily and (b) monthly versus the number of tourists (lines) each month from 1 April to 30 November of 2018 (light grey) and 2019 (dark grey). Horizontal bars below the chart show the period when residents and migratory breeding females and males are present.

### Unique turtles identified from Instagram records

In both years, about half of all entries were of sufficient quality to be compared against the photoidentification database (49% and 59% respectively). Identification was more successful for underwater entries (66 to 78% for 2018 and 2019) as compared to boat entries (41 and 49%) (Supplementary table 2).

In 2018 and 2019, 139 and 122 unique individuals were identified, respectively, of which 40 and 37 were residents (Supplementary table 2). While residents represented just *∼*30% of all identified individuals, there was a strong observation bias for this group (82% of entries in 2018 and 88% in 2019; Supplementary table 2).

Overall, 86 and 91% of boat entries, and 66% and 81% of underwater, in 2018 and 2019 were of resident turtles, respectively. Interestingly, the residents that were heavily targeted by boats (5 mature males and 1 unknown; about 70% of entries in both years) in both years had far fewer underwater entries, with other residents being targeted underwater, particularly in 2019 (Figure 4). For instance, four residents (all immature) accounted for 56% of underwater entries in 2019. Of note, 44% of residents were recorded in both years. Detailed number of entries per individual turtle can be found in Supplementary figure 2.

**Figure 4.**
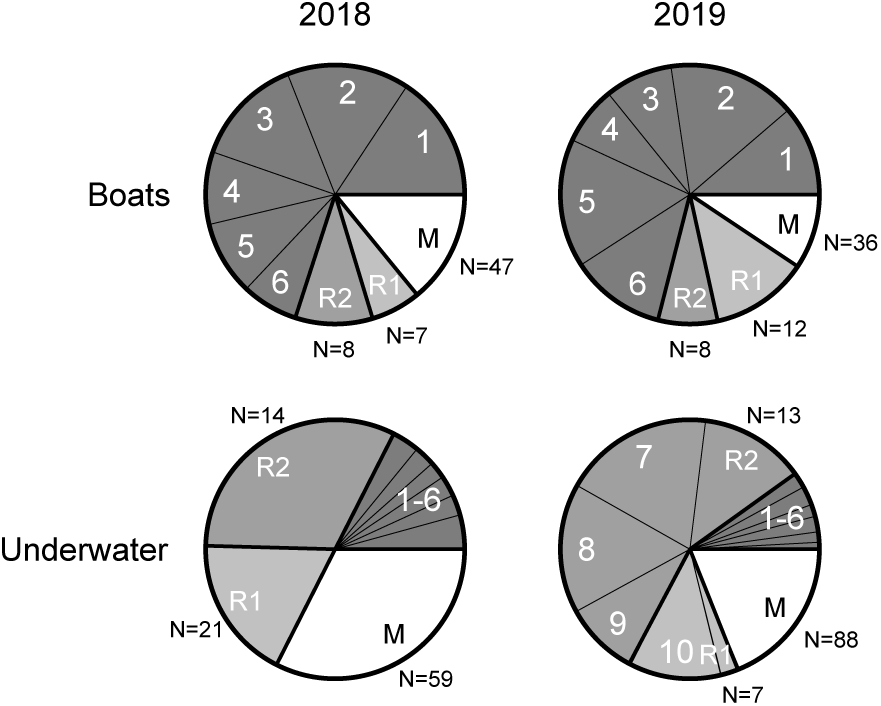
Relative number of entries of uniquely identified resident (light to dark grey shading) and migrant (white shading) individuals from boat and underwater sightings in 2018 (n = 497 and 234 entries) and 2019 (n = 673 and 464 entries). 1-6 (dark grey) = six uniquely identified resident individuals sighted in both years that represented a large portion of boat entries; 7-10 = other individuals of note representing a large portion of underwater entries in 2019; R2 (intermediate grey) = other residents sighted in both years; R1 (light grey) = residents sighted in one of the two years. N for R2, R1, and M represents the number of unique individuals within that group.

### Temporal variation in tourism pressure on residents

In both years, even when breeding individuals were present in June and July, resident turtles represented 60-70% of entries, rising up to 100% of entries in August onwards, after breeding individuals had departed (Figure 5).

**Figure 5.**
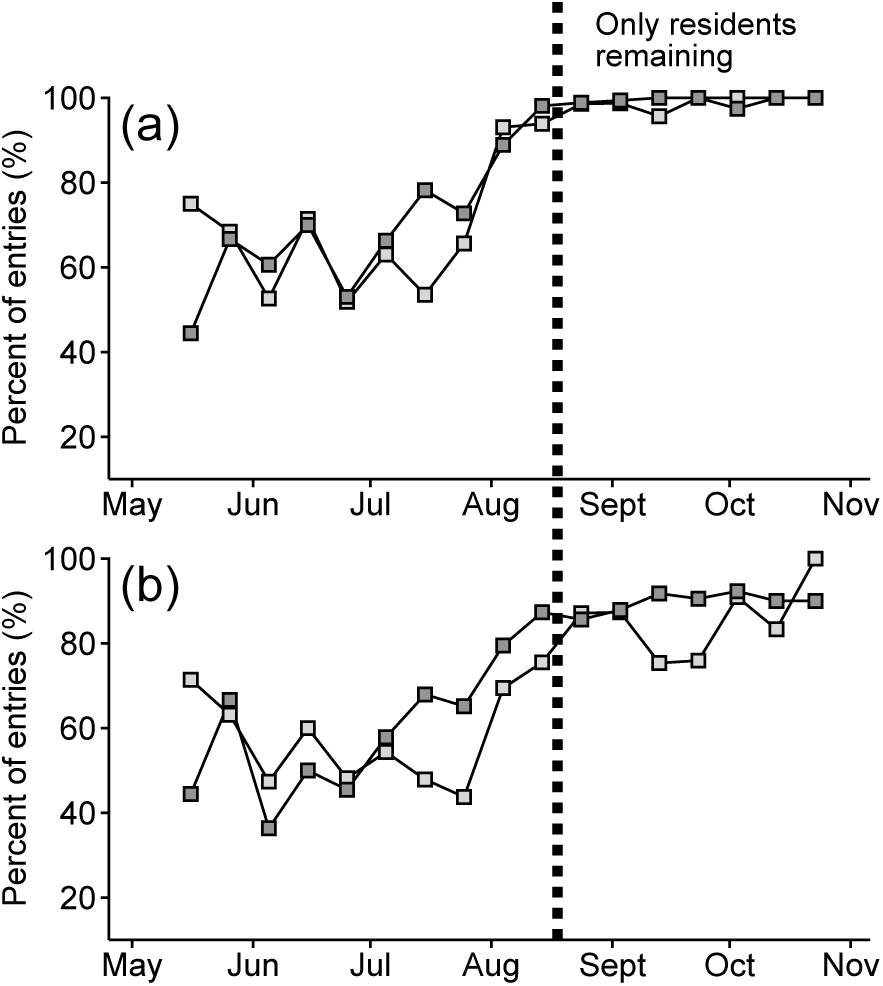
Temporal variation in the percentage of (a) all residents and (b) the top 10 residents representing social media entries out of all entries in 2018 (light grey squares) and 2019 (dark grey squares). Percentages were grouped for 10-day periods with each square placed in the middle of these periods. A 10 day period was chosen because it was large enough smooth wild oscillations and small enough to capture temporal changes. Data before 11 May and after 28 October were not plotted due to the very small number of entries in these periods.

### Risk of trauma

Overall, out of all 54 residents observed in 2018 and 2019, 17% (n = 9) had evidence of carapace damage by boats. Of these, two individuals (both males) were only observed in 2019, and age of the injury could not be determined. The other seven individuals, of which five were males, were among the 24 residents documented in both years, all of which had evidence of propeller injury on their carapace. Of these seven individuals, three sustained new damage within the 2019 season, of which one was fatal (ARCHELON unpublished data). Another turtle sustained new damage at some point between 2018 and 2019, one had been damaged within the 2016 season, and the other two were already injured when first observed (2016 and 2017, respectively) (Figure 6). Of note, four of these seven individuals (including the fatality) were among the top 10 residents observed by boats in both 2018 and 2019 (Figure 4; Supplementary figure 4).

**Figure 6.**
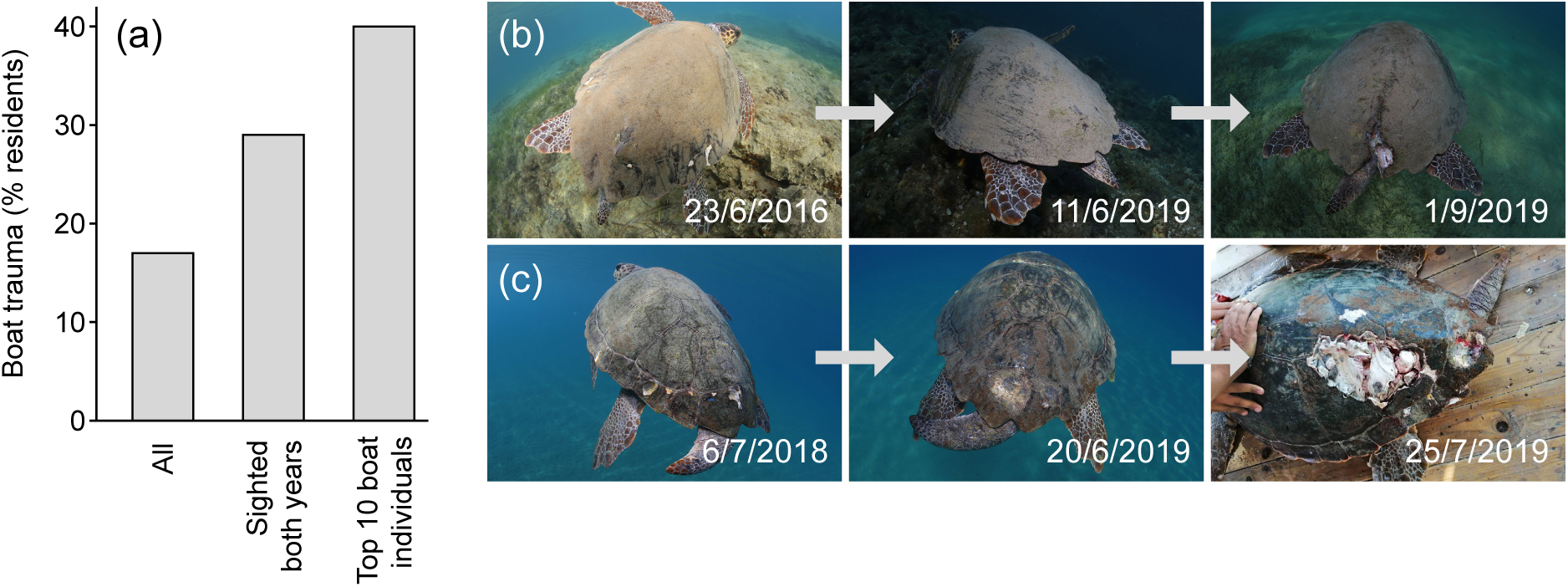
Boat trauma (percentage) recorded for (a) all 54 documented residents recorded across the two survey years (n = 9), the 24 residents recorded in both years (n = 7), and the 10 most frequently sighted residents by boats (n = 4). Examples of boat trauma sustained by two residents within the 2019 season: (b) t235, a male showing no trauma in 2016 and June 2019, but boat trauma sustained by September 2019; (c) t110, (“1” in Figure 4) a male showing no trauma in 2018, a healed trauma in June 2019 (sustained at some point between 2018 and 2019) and fatal trauma sustained in July 2019. This individual died in September 2019 after being sent for treatment at the ARCHELON rescue center, Athens. Photograph credits: Kostas Papafitsoros apart bottom right photograph (Giorgos Rallis, used with permission).

## Discussion

By combining social media with photo-identification, our study confirmed that resident male and immature turtles are subject to disproportionately higher levels of viewing both by boats and in-water observers (swimmers/snorkelers). In particular, this small group of up to 40 individuals accounted for over 60% of total viewings, even when the entire seasonal breeding population of over 200 individuals was present. During both seasons, six resident individuals were primarily targeted during boat observations, potentially reflecting their high fidelity to particular locations and indicating low disturbance by this activity. However, there is strong evidence of very high risk of trauma or mortality to these individuals, demonstrating the importance of ensuring the national park authority and local turtle viewing operators are made aware of their disproportionate use of resident turtles, and the need to develop ways to ensure this major economic resource is used wisely.

The high viewing pressure on resident turtles indicates that operators are targeting a specific area of the bay where these individuals are more likely to be found, forgoing random searching for turtles, to guarantee viewings throughout the season.This supports the findings of Schofield et al. (2015), who demonstrated that most boat-based watching activity was concentrated in the vicinity of Agios Sostis to Laganas Village (Figure 1). Thus, even when migrant breeders are present in the bay in large numbers, operators preferentially target areas used by residents rather than randomly searching for turtles. Previous telemetry studies (Dujon et al., 2018) and long-term direct in-water observations confirm high utilization of this part of the bay by residents (Papafitsoros and Schofield, 2016). Fidelity to foraging sites has been widely recorded in sea turtle species (Limpus et al., 1992; Broderick et al., 2007; Rees et al., 2013; Shimada et al., 2020). Other species also exhibit high fidelity to certain foraging or breeding locations (e.g. rays, Barr and Abelson, 2019; sharks, Cisneros-Montemayor et al., 2013; marine mammals, Christiansen and Lusseau, 2014), with these studies demonstrating that these sites are targeted by operators to guarantee viewings. Consequently, bias in viewing locations likely leads to greater pressure on certain individuals or groups in different periods. The current study, however, demonstrated that bias in viewing resident turtles remained high, even when the population was inflated by the presence of migrants.

However, the consistently high number of sightings of the same six turtles by boats in both years and the four turtles by swimmers/snorkelers in 2019 suggests that high viewing pressure from boats or swimmers/snorkelers does not seem to disrupt these individuals as they appeared to remain in the same area. Viewing pressure on wildlife may lead to disturbed individuals shifting to alternative sites (Semeniuk et al., 2009; Williams et al., 2009). In other cases, increased human disturbance has low impact (e.g. marine mammals New et al., 2013; Holcomb et al., 2009). Obtaining information on animals targeted by swimmers/snorkelers who are not part of organized groups is logistically challenging (e.g. marine mammals, King and Heinen, 2004; Dans et al., 2017, sea turtles, Griffin et al., 2017) but social media here provided an opportunity to explore the impact of this phenomenon. The extent of impact is considered to be strongly linked to the habitat and behavior of the animals using it (Gill et al., 2001; Williams et al., 2009). High viewing pressure might trigger avoidance behavior and, thus, impact the energy expenditure of individuals or their ability to accumulate energetic reserves, such as foraging efficiency or energy conservation strategies (Christiansen et al., 2013; Fossette et al., 2012). This is a potential issue in our case, as the resident turtles were foraging on molluscs submerged in sandbanks (representing residents, typically adult males, primarily targeted by boats) or on sponges in reefs immediately adjacent to shore (representing residents, typically immature turtles, primarily targeted by swimmers/snorkelers) (Kostas Papafitsoros and Gail Schofield, personal observations). In particular, while underwater viewings were spread among a large number of individuals in 2018, this changed in 2019, which might have been due to a key foraging spot being made accessible, altering the dynamic of underwater viewings (Figure 1 and Supplementary figure 3). This demonstrates that social media could be used to identify shifts in viewing pressure facilitating evidence based management of this activity (Sutherland et al., 2004).

In addition to swimming/snorkeling with animals potentially impacting the ability of animals to build sufficient energy reserves, high and repeated exposure to boats could increase the risk of trauma or mortality to individuals through propeller and boat strikes, as documented for sea turtles (Arianoutsou, 1988; Denkinger et al., 2013; Lester et al., 2013), and other species (e.g. bottlenose dolphins, Wells and Scott, 1997, manattees, Calleson and Frohlich, 2007). It was recently shown that adult males that breed on Zakynthos have a much lower annual survival rate than females, and that males tend to use habitats directly adjacent to shore when foraging (Schofield et al., 2020). These suggestions based on Fastloc GPS tracking data are directly supported by the social media entries assimilated in this study, showing the nearshore use of adult male turtles plus the increased risk of boat strike. Based on the first six years of the photo-identification database (2000 to 2006), (Schofield et al., 2013) estimated that over 40% of turtles frequenting the NMPZ marine area had sustained some sort of physical injury from human activities (boat strike and fishing gear). The current study confirms propeller strikes represent a major threat to the resident turtles; however, it was not possible to determine whether these injuries arise during organised wildlife watching activities, by private hired boats operated by tourists, or other unidentified cause (e.g. Illegal fishing boat activity).

The high risk of trauma or mortality of turtles due to propeller or boat strike should be of concern to operators (Keough and Blahna, 2006; Duprey et al., 2008; Cisneros-Montemayor et al., 2013), particularly as this industry is dependent on viewing residents during the peak tourist season from August onwards when most migrants departed, which was clearly shown by the social media entries. Operators tend to comply with existing guidelines only when sufficient numbers of turtles are present (Schofield et al., 2015). Our results allow us to quantify the value of viewed turtles to operators, which might alter their interest in complying to regulations. Specifically, resident turtles represent 80% of the annual viewing revenue which is estimated at more than 2.7 million euros. Of note is the fact that with 70% of viewings being attributed to the same six residents in both years, each resident essentially contributes an estimated 315,000 euros per year (six month viewing period) to the local economy, which is significant amount when compared to the minimum Greek wage over the same 5-month period (3500 euros). Without question, preventing human-induced mortality of male turtles should be a management focus, particularly in light of the known lower survival rates of males in this population (Schofield et al., 2020), the lower relative numbers of adult males present in the breeding population (1 male to four females; Schofield et al., 2017) and the fact that offspring production is strongly female biased globally, with that bias possibly persisting in the adult populations (Hays et al., 2014). Wildlife watching vessels have been shown to primarily use a restricted area of the bay (Schofield et al., 2015), which overlaps with the area frequented by resident male and immature turtles for foraging (based on Fastloc GPS tracking data; Dujon et al., 2018), particularly the key six individuals (Kostas Papafitsoros-personal observations).

An important management intervention could include the compulsory use of propeller guards on all viewing and private vessels used within Laganas Bay, which combined with a strong enforcement of the current 6 knot limit in maritime zones B and C, could significantly reduce propeller strikes in this focal area. Due to the strong overlap in operator area use with resident turtle area use (Schofield et al., 2013, Schofield et al., 2015, Dujon et al., 2018), another possible solution could be to restrict the use of this part of the bay (currently the least restricted maritime zone C or key foraging spots; Figure 1), with a temporary (time of day or day of month) no-entry “refuge” zone for residents (Williams et al., 2009). The restriction of unregulated recreational craft has been shown to have a positive impact on killer whales and dolphins (Jelinski et al., 2002; Duprey et al., 2008). However, on the flipside, encouraging operators and other vessels to change area use, would lead to increased viewings of reproductive females. Our results showed that boat and underwater viewing pressure on reproductive females is currently relatively low, corroborating previous inferences (Schofield et al., 2015). However, the energetic cost of viewing females, even at low intensities, might be disproportionately high (e.g. as documented for cetaceans by Christiansen and Lusseau, 2014), due to the impacts on energetic resources and hence reproductive capacity (Fossette et al., 2012). Consequently, such a measure involves a certain risk, since we do not know whether females would move to alternative areas when subjected to intense viewings, and whether alternative areas have sufficient resources sought by females.

## Conclusion

This study validates the utility of large quantities of social media data for capturing data on individuals at the population level and how this varies over time. Through using social media data, we confirmed that viewing pressure is disproportionately biased towards very small numbers of resident turtles, even when large numbers of breeding individuals are present. This bias likely occurs for other taxa where viewing encompasses mixed resident and migrant populations; however, due to their different habitat needs, operators probably target sites where individuals are consistently present across a season (i.e. those used by residents). Thus, social media represents a useful approach towards distinguishing bias in wildlife viewing and changes to this bias, providing a means to quantify the economic value of targeted individuals to operators, providing a strong incentive to comply and even contributing towards strengthening regulations.

## Acknowledgments

The authors thank George Valais for logistical support and Ioannis Giovos for useful discussions. The first author acknowledges the support of the Oceanic Society through a SWOT small grant 2017.

## Supplementary material

List of hashtag code names as well as the locations that were used for the Instagram search:

### Hashtags

- #zakynthos
- #zakynthosisland
- #zakyntos
- #zakintos
- #zakinthos
- #zakhyntos
- #zacinto
- #zacintos
- #zante
- #zante2018
- #zante2019
- #zakynthos2018
- #zakynthos2019
- #zakynthos_greece
- #zante_greece
- #laganas
- #kalamaki
- #laganas
- #gerakas
- #laganas
- #dafni
- #laganasbeach
- #kalamakibeach
- #gerakasbeach
- #dafnibeach
- #agiossostis
- #cameoisland
- #cameo
- #marathonisi
- #marathonissi
- #loggerhead
- #loggerheadturtle
- #loggerheadseaturtle
- #caretta
- #carettacaretta
- #careta
- #caretacareta
- #karetakareta
- #kareta
- #zolwie
- #zolw
- #zelva
- #tartaruga
- #turtlespotting

### Locations

- Zákynthos, Zakinthos, Greece
- Zante, Zakinthos, Greece
- Zakynthos Island
- Marathonisi Turtle Island Zante
- Marathonisi, Zakinthos
- Laganas, Zakynthos, Greece
- Laganas Beach
- Laganas Beach, Turtle Spotting
- Laganas
- Kalamáki, Zakinthos, Greece
- Kalamaki Beach
- Gerakas beach, Zakynthos
- Agios Sostis, Zakynthos
- Cameo Island, Zante, Greece
- Cameo Island Club
- Vasilikόs, Zakinthos, Greece
- Turtle Trip

---

**Supplementary figure 1.**
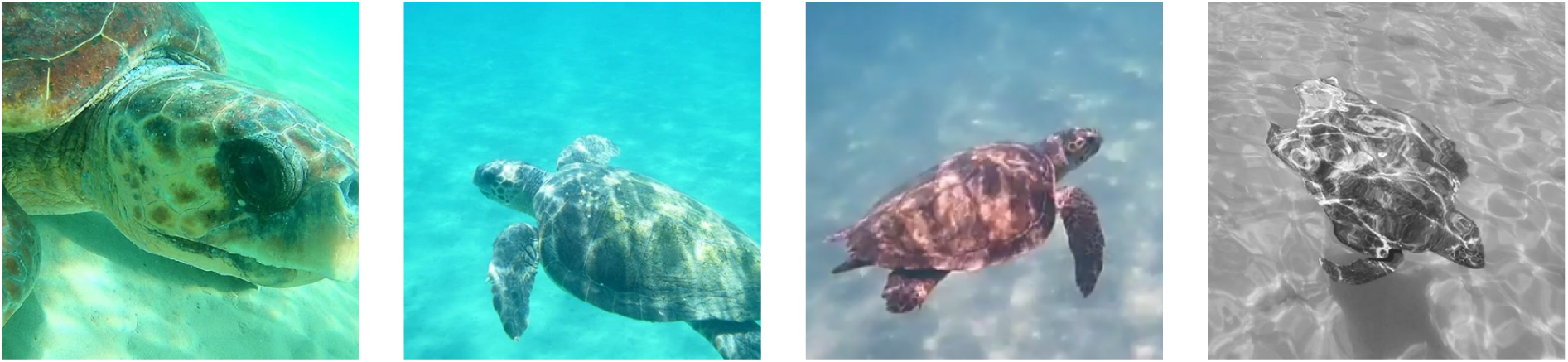
Example of inspected Instagram photos: (a) Example of photograph that was of good enough quality for photo-identification (b) Example of photograph of medium quality but still sufficient for photo-identification (c) Example of photograph of insufficient quality for photo-identification but still possible to be identified as a male (d) Example of a photograph of insufficient quality to identify individual or sex (Note: many times males hide tails under the carapace). Photograph credits: (a) Giannis Xenos, (b-c) Rémi Rossignol, (d) Margarita Koulouri, used with permission.

**Supplementary table 1.**
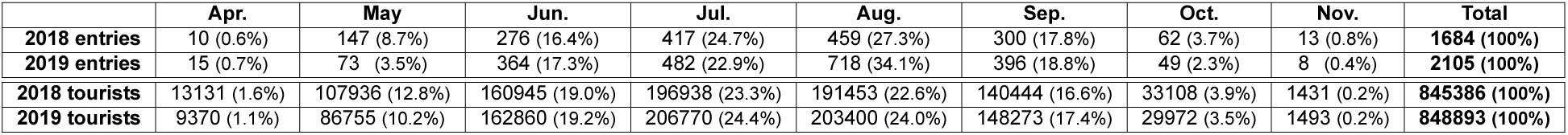
Monthly number and percentages of inspected Instagram entries and tourists for 2018 and 2019. Tourist numbers represent the monthly number of air passenger arrivals according to the Hellenic Civil Aviation Authority (CAA) (http://www.ypa.gr).

**Supplementary table 2.** Numbers and percentages of boat, underwater and land entries for 2018 and 2019.Each column is further broken down to non-identified and identified entries, and whether individuals were non-residents, old (existing) residents and (potentially) newly detected residents. The last column contains numbers and percentages of the identified individuals for both years. Percentages in red and green are considered row and column-wise respectively.

| 2018 | Boat entries | Underwater entries | Land entries | Total entries | Identified Individuals |
| --- | --- | --- | --- | --- | --- |
| Non-identified | 727 (59.4%) | 119 (33.7%) | 8 (7.5%) | 854 (50.7%) |  |
| Identified (IDed) | 497 (40.6%) | 234 (66.3%) | 99 (92.5%) | 830 (49.3%) | 139 (100%) |
| Non-residents | 70 (14.1% of IDed) | 76 (32.5% of IDed) | 0 (0.0% of IDed) | 150 (18.1% of IDed) | 99 (71.2%) |
| Old residents | 354 (71.2% of IDed) | 110 (47.0% of IDed) | 92 (92.9% of IDed) | 556 (67.0% of IDed) | 22 (15.8%) |
| New residents | 73 (14.7% of IDed) | 48 (20.5% of IDed) | 7 (7.1% of IDed) | 124 (14.9% of IDed) | 18 (13.0%) |
| Total | 1224 (100%) (72.7%) | 353 (100%) (20.9%) | 107 (100%) (6.4%) | 1684 (100%) (100%) |  |
| 2019 | Boat entries | Underwater entries | Land entries | Total entries | Identified Individuals |
| Non-identified | 714 (51.5%) | 134 (22.4%) | 11 (9.2%) | 859 (40.8%) |  |
| Identified (IDed) | 673 (48.5%) | 464 (77.6%) | 109 (90.8%) | 1246 (59.2%) | 122 (100%) |
| Non-residents | 63 (9.4% of IDed) | 88 (19.0% of IDed) | 0 (0.0% of IDed) | 151 (12.1% of IDed) | 85 (69.7%) |
| Old residents | 571 (84.8% of IDed) | 312 (67.2% of IDed) | 109 (100.0% of IDed) | 992 (79.6% of IDed) | 24 (19.7%) |
| New residents | 39 (5.8% of IDed) | 64 (13.8% of IDed) | 0 (0.0% of IDed) | 103 (8.3% of IDed) | 13 (10.6%) |
| Total | 1387 (100%) (65.9%) | 598 (100%) (28.4%) | 120 (100%) (5.7%) | 2105 (100%) (100%) |  |

**Supplementary figure 2.**
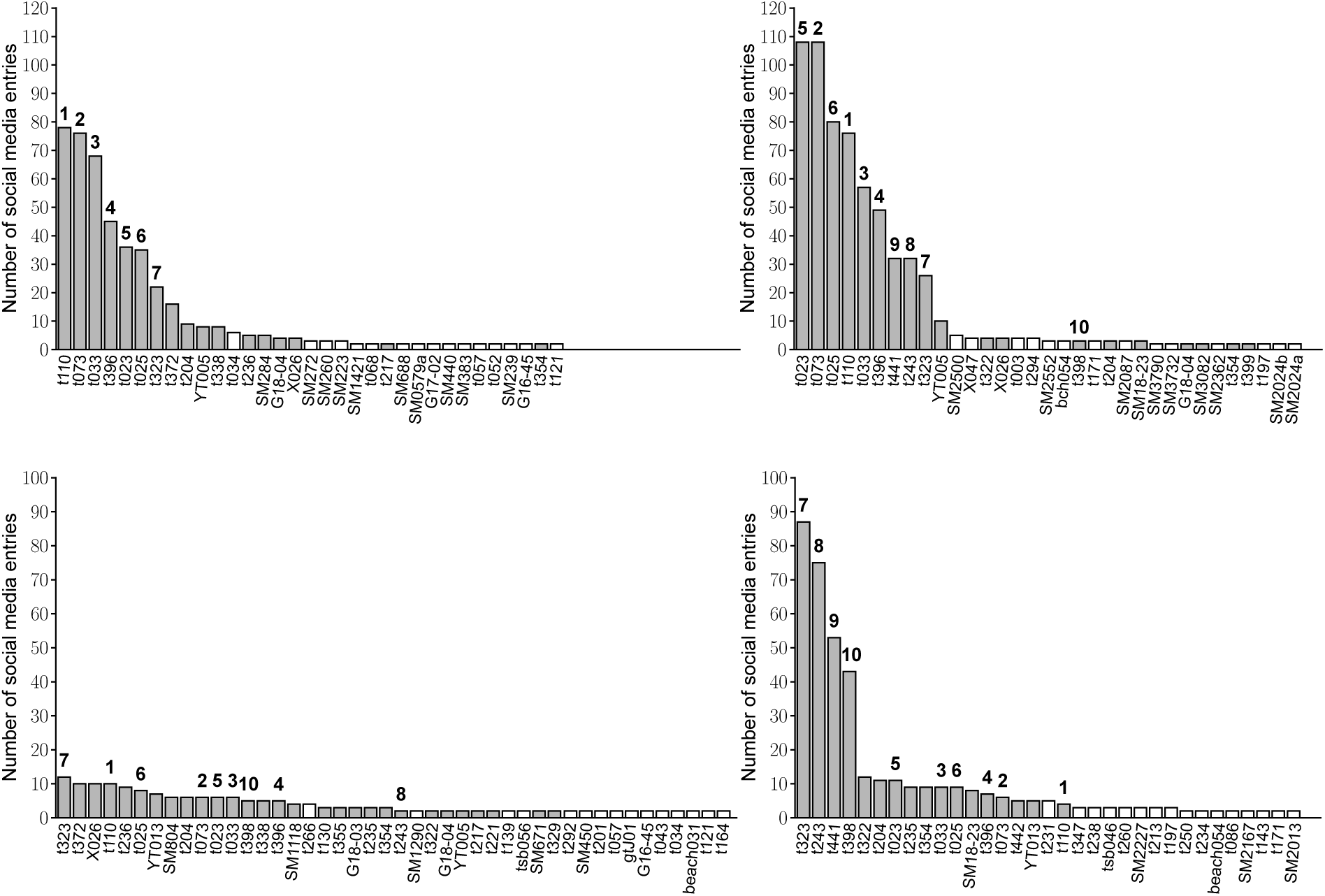
Number of Instagram entries per turtle for boats (top) and underwater (bottom) entries for 2018 (left) and 2019 (right). Grey and white bars denote resident and non-resident individuals respectively. The x-axis labels displays the database code names of the uniquely identified individuals while the bold numbers (1-10) correspond to the coding of the 10 individuals in Figure 4. Only individuals with more than 1 entry are shown.

**Supplementary figure 3.**
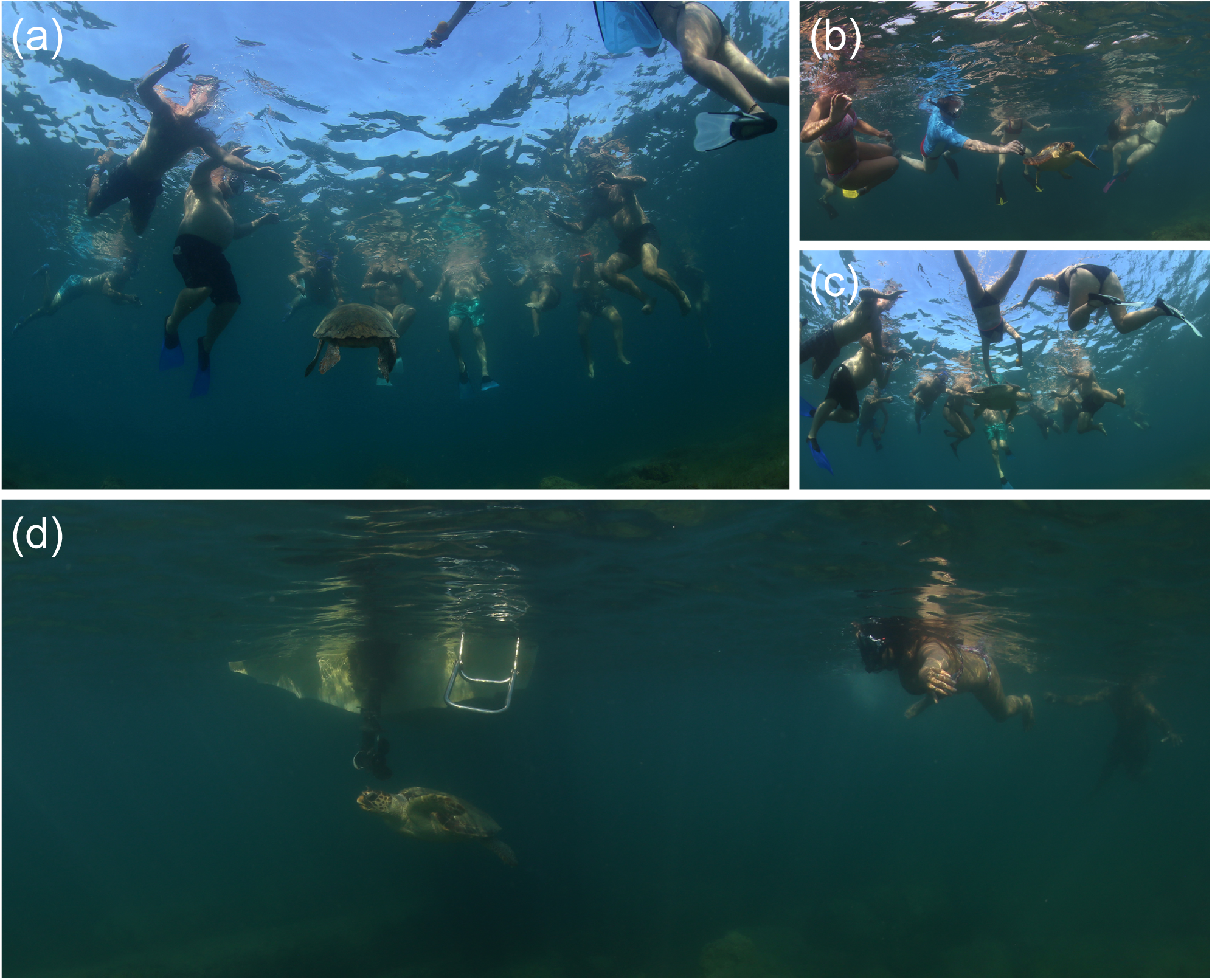
(a-c) Documentation of the intense swimmer pressure on the foraging spot of Agios Sostis (Figure 1(b)) during August-September 2019, frequently used by the four above individuals “t323”, “t243”, “t441”, “t398” (coded with “7”, “8”, “9”, “10” in Figure 4). (a) & (c): individual “t243” on 5 September 2019. (b): individual “t398” on 3 September 2019. (d): Individual “t323”, swimming next to a boat propeller under simultaneous viewing by swimmers and boat, 20 August 2018. Photograph credits: Kostas Papafitsoros.

**Supplementary figure 4.**
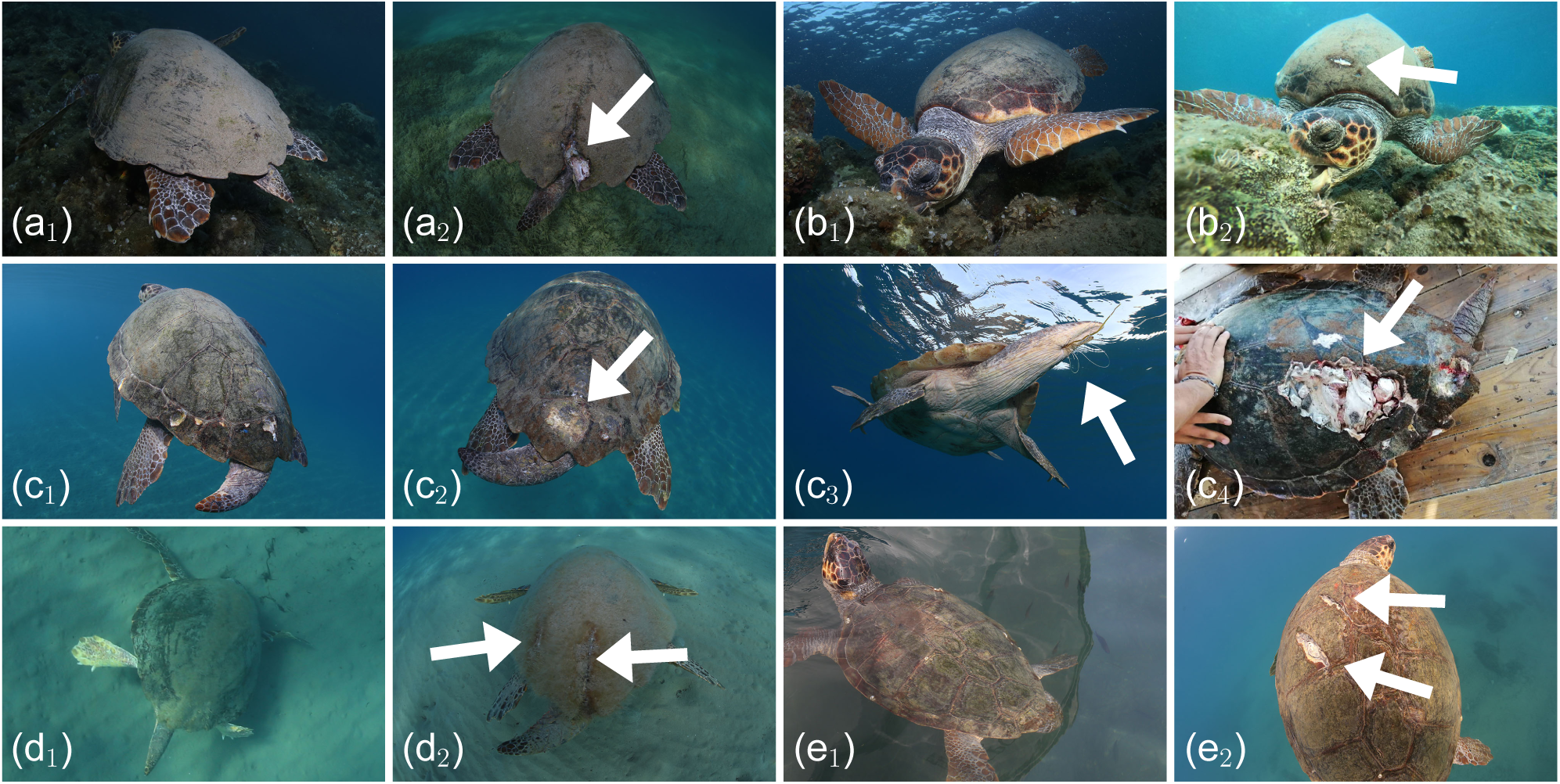
Documentation of recent boat injuries to resident turtles of Laganas Bay: (a_1_-a_2_) Resident male “t235” on 11 June 2019 (a_1_) and 1 September 2019 (a_2_). (b_1_-b_2_) Resident immature turtle (“8” in Figure 4) on 5 September 2019 (b_1_) and end of September 2019 (b_2_). Note: this individual was 8^*th*^ in the boat viewings in 2019, see top left bar plot in Supplementary figure 2. (c_1_-c_4_) Resident male (“1” in Figure 4), 6 July 2018 (c_1_), 20 June 2019 (c_2_), 19 June 2019 (c_3_) (note the fishing line coming out from the cloaca) and 26 July 2019 (c_4_). Note: this individual was 1^*st*^ and 4^*th*^ in boat viewings in 2018 and 2019 respectively (1^*st*^ in 2019 as well when considering viewings only till 26 July 2019). The animal eventually died in ARCHELON rescue center, September 2019. (d_1_-d_2_) Resident male “X026” on 23 June 2018 (d_1_) and 16 June 2019 (d_2_). (e_1_-e_2_) Resident male (“3” in Figure 4) on 23 June 2016 (e_1_) and 2 October 2016 (e_2_). Note: this individual was 3^*rd*^ and 5^*th*^ in the boat viewings in 2018 and 2019 respectively. Photograph credits: Kostas Papafitsoros apart from (b_2_) Lisa Matthiesen and (c_4_) Giorgos Rallis, both used with permission.

**Supplementary figure 5.**
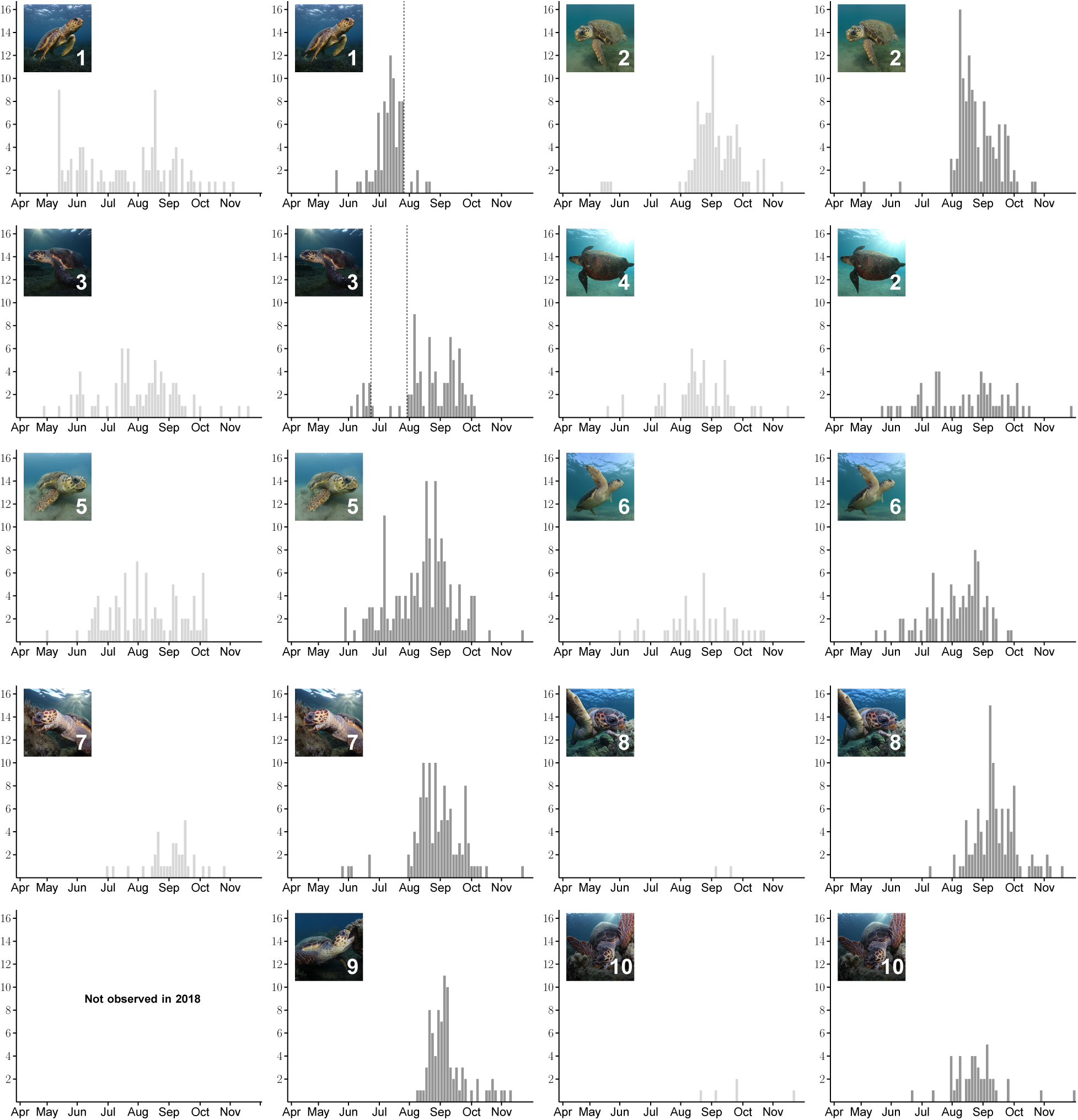
Number of Instagram entries for the top 10 individuals “1”-”10” of Figure 4 for 2018 (light grey) and 2019 (dark grey). All bars denote number of entries in 3 days interval. **1**. Database code “t110”, resident male: First in boat entries in 2018, and very intensely observed in 2018 and 2019 until 26 July 2019 (dashed line), when it was fatally injured by a propeller possibly during turtle spotting activities. **2**. Database code “t073”, resident, sex not determined: Similar entry distribution pattern in both years, with very few entries at the end of May, no entries in the following two months, with a sharp increase from August onwards at the end of the nesting season. **3**. Database code “t033”, resident male: Similar temporally uniform pattern in both years. Notice the gap between 22 June 2019-27 July 2019 (dashed lines), in which it was absent from the area. The turtle was transferred to ARCHELON rescue center with a hook and was released from Athens on 3 July 2019, returning to Zakynthos in just over 3 weeks. **4**. Database code “t396”, resident male: Uniform entry distribution pattern in both years. **5**. Database code “t023”, resident male: Uniform entry distribution pattern in both years. **6**. Database code “t025”, resident male: Uniform entry distribution pattern in both years. **7-10**. Database codes “t323”, “t243”, “t441”, “t398”, all resident immature turtles: Number of entries increased in August-September 2019, possibly due to parallel increase in tourist numbers using the same area, see also Supplementary figure 3. Individual “t441” was not observed in 2018. Photograph credits: Kostas Papafitsoros

